# The Hunchback temporal transcription factor determines motor neuron axon and dendrite targeting in *Drosophila*

**DOI:** 10.1101/554188

**Authors:** Austin Q. Seroka, Chris Q. Doe

## Abstract

The generation of neuronal diversity is essential for circuit formation and behavior. Morphological differences in sequentially born neurons could be due to intrinsic molecular identity specified by temporal transcription factors (henceforth called intrinsic temporal identity) or due to changing extrinsic cues. Here we use the *Drosophila* NB7-1 lineage to address this question. NB7-1 sequentially generates the U1-U5 motor neurons; each has a distinct intrinsic temporal identity due to inheritance of a different temporal transcription factor at time of birth. Here we show that the U1-U5 neurons project axons sequentially, followed by sequential dendrite extension. We misexpress the earliest temporal transcription factor, Hunchback, to create “ectopic” U1 neurons with an early intrinsic temporal identity but later birth-order. These ectopic U1 neurons have axon muscle targeting and dendrite neuropil targeting consistent with U1 intrinsic temporal identity, rather than their time of birth or differentiation. We conclude that intrinsic temporal identity plays a major role in establishing both motor axon muscle targeting and dendritic arbor targeting, which are required for proper motor circuit development.

## Introduction

Axon and dendrite targeting is an essential step in neural circuit formation, and may even be sufficient for proper connectivity in some cases, as postulated in Peters’ Rule (Peters and Feldman, 1976; Rees et al., 2017; Stepanyants and Chklovskii, 2005). In both *Drosophila* and mammals, individual progenitors generate a series of neurons that differ in axon and dendrite targeting (Doe, 2017; Kohwi and Doe, 2013; Li et al., 2013a; Pearson and Doe, 2004; Rossi et al., 2016). In all examples, neurons born at different times have intrinsic molecular differences due to temporal transcription factors (TTFs) present at their time of birth (reviewed in Kohwi and Doe, 2013), which could specify neuronal morphology. Conversely, there are likely changing extrinsic cues present at the time of neuronal differentiation that could also influence neuronal morphology, such as modulation of global pathfinding cues or addition of axon and dendrite processes throughout neurogenesis. Teasing out the relative contributions of intrinsic or extrinsic factors requires heterochronic experiments where either intrinsic or extrinsic cues are altered to create a mismatch, and the effects on axon and dendrite targeting are assessed.

Several experiments highlight the importance of extrinsic cues present at the time of neuronal differentiation in establishing axon or dendrite targeting. For example, transplantation of rat fetal occipital cortical tissue into the rostral cortex of a more developmentally mature newborn host results in axonal projections characteristic of the host site (O’Leary and Stanfield, 1989; Schlaggar and O’Leary, 1991; Stanfield and O’Leary, 1985). Similarly, transplantation of embryonic day 15 fetal occipital tissue into newborn occipital cortex reveals that the transplanted tissue receives thalamic projections typical of the host site and developmental stage (Chang et al., 1986). More recent work in zebrafish shows that vagus motor neurons extend axons sequentially to form a topographic map, and that the time of axon outgrowth directs axon target selection (Barsh et al. 2017). In all of these experiments, there are unlikely to be changes in intrinsic temporal identity of the donor neurons, suggesting that the heterochronic neurons are establishing neuronal morphology in response to different environmental cues present at their time of differentiation.

In contrast, heterochronic experiments where donor neurons maintain donor identity are more consistent with intrinsic temporal identity specifying neuronal axon and dendrite targeting. For example, heterochronic experiments in ferrets show that late cortical progenitors transplanted into younger hosts generate neurons with late-born deep layer position and subcortical axonal projections (McConnell, 1988). Transplantation of older post-natal cerebellum into embryonic host mice results in the neurons maintaining donor “late-born” identity based on molecular markers and neuronal morphology (Jankovski et al., 1996). Similarly, experiments done in grasshopper embryos show that delaying the birth of the first-born aCC motor neuron in the NB1-1 lineage leads to defects in the initial axon trajectory (extending anterior instead of posterior) but the temporally delayed aCC invariably finds and exits through the proper nerve root in the adjacent anterior segment (Doe et al., 1986). In all of these heterochronic experiments, it is likely that intrinsic temporal identity is unaltered and helps maintain donor neuron identity despite their altered time of differentiation. However, none of these experiments show that intrinsic temporal identity is unchanged in the transplanted neurons, and none of these experiments manipulates intrinsic temporal identity to directly assess its role in establishing proper axon or dendrite targeting.

We sought to test the relative contribution of neuronal time of differentiation versus neuronal intrinsic temporal identity in establishing motor neuron axon and dendrite targeting. Our model system is the NB7-1 lineage in the *Drosophila* ventral nerve cord (VNC), a segmentally repeated structure analogous to the mammalian spinal cord. The VNC offers several benefits to the study of neurogenesis due to its individually identifiable neuroblasts (NBs) which produce a stereotyped sequence of distinct neuronal cell types whose identities are determined by a well-characterized temporal transcription factor (TTF) cascade (reviewed in Doe, 2017; Kao and Lee, 2010; Kohwi and Doe, 2013; Rossi et al., 2016; Skeath and Thor, 2003) (Fig. 1; Supplemental Fig. 1). For example, NB7-1 sequentially expresses the four TTFs Hunchback (Hb), Kruppel, Pdm, and Castor. During each NB TTF expression window a different motor neuron is born: U1 and U2 during the Hb window; U3, U4, and U5 during the later three TTF windows (Isshiki et al., 2001; Kanai et al., 2005; Kohwi et al., 2013; Pearson and Doe, 2003). Importantly, the two Hb+ U1-U2 motor neurons have a morphology, neuropil targeting, and connectivity that is clearly different from the later-born U3-U5 motor neurons (Fig. 1; Supplemental Fig. 1). The ability to individually identify the U1-U5 neurons, and to cleanly change their intrinsic temporal identity in an otherwise normal CNS, make the NB7-1 lineage an ideal system to study the relative contribution of time of differentiation and intrinsic temporal identity for establishing neuron morphology, targeting, and connectivity.

**Fig. 1.**
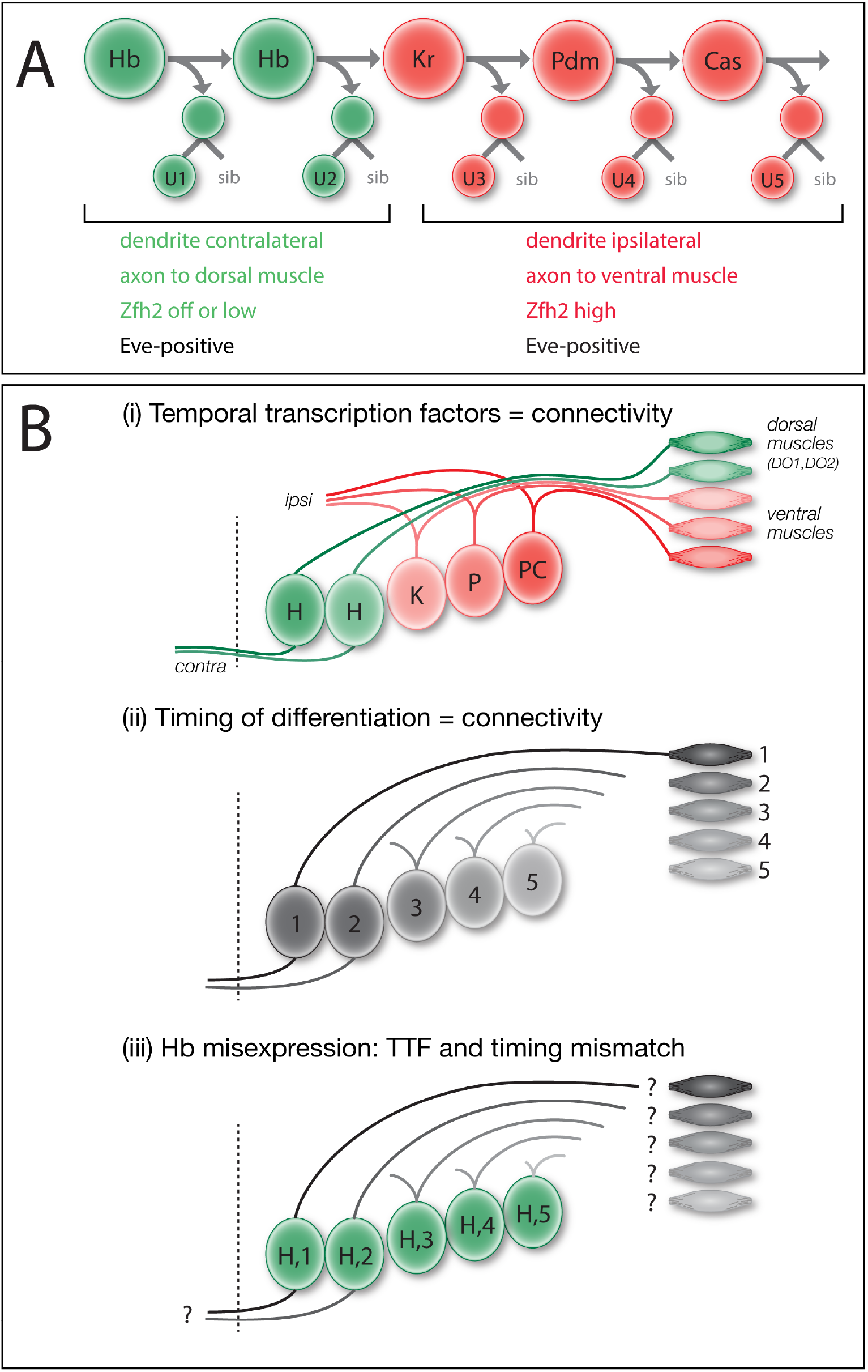
Models: intrinsic temporal identity or time of differentiation determines U1-U5 motor neuron morphology. (A) NB7-1 (top row) sequentially expresses the temporal transcription factors Hb, Kr, Pdm, and Cas. The U1-U2 neurons (bottom row) born during the Hb window have an “early-born” identity (green) characterized by contralateral dendrites, an axon projection to dorsal body wall muscles DO1 and DO2, and little or no nuclear Zfh2. The U3-U5 neurons born after the Hb window have a “late-born” identity (red) characterized by ipsilateral dendrites, an axon projection to more ventral muscles DA3/LL1 and have high nuclear Zfh2. All U1-U5 neurons have nuclear Eve. (B) Models for specification of U1-U5 axon and dendrite targeting. (i) Intrinsic temporal identity could determine axon and dendrite targeting. (ii) Neuronal time of differentiation could determine axon and dendrite targeting. (iii) Misexpression of Hb can generate late-differentiating neurons with an early intrinsic temporal identity; this mismatch reveals which mechanism is more important for axon and dendrite targeting.

Previously we showed that misexpression of Hb throughout the NB7-1 lineage results in an extended series of “ectopic U1” motor neurons based on molecular markers and axon projections to dorsal body wall muscles (Isshiki et al., 2001; Pearson and Doe, 2003); however, the ectopic U1 motor neurons were not assayed for their specific muscle targets, nor was dendrite morphology and targeting assessed, nor was it known if U1-U5 motor neurons extended axons or dendrites synchronously or sequentially. Here we focus on difference between early-born Hb+ U1-U2 neurons and later-born U3-U5 neurons. U1-U2 are bipolar, have contralateral dendrites, and innervate dorsal body wall muscles; in contrast, U3-U5 neurons are monopolar, have ipsilateral dendrites, and innervate more ventral body wall muscles. Although there are molecular differences between U1-U2 and between U3-U5 (Isshiki et al., 2001), in this paper we focus on the major morphological differences between these two groups of neurons. We show for the first time that the U1-U5 neurons extend axons sequentially, and subsequently extend dendrites sequentially. Here we test whether U1-U5 motor neurons project to their normal CNS and muscle targets due to their intrinsic temporal identity (Fig. 1Bi) or due to their time of differentiation (Fig. 1Bii) – two mechanisms that are normally tightly correlated. To break this correlation, we misexpress the early TTF Hb specifically in the NB7-1 lineage to create “ectopic U1” motor neurons with an early intrinsic temporal identity but late time of differentiation (Fig. 1Biii). Moreover, we show that the heterochronic placement of an “ectopic U1” into the later developmental environment does not affect the ability of the “ectopic U1” to project dendrites to the proper neuropil domain or axons to the proper body wall muscle. Our results show that intrinsic temporal identity is an important determinant of neuronal morphology and axon and dendrite targeting.

## Results

### U1-U5 motor neurons extend axons and dendrites sequentially

To determine whether U1-U5 motor neuron axon target selection is correlated with intrinsic temporal identity or time of differentiation, we first needed to investigate the timing of U1-U5 motor neuron axon outgrowth. If the U1-U5 motor neurons have synchronous axon outgrowth, despite being born sequentially, we can rule out time of axon outgrowth as a mechanism for specifying their differential axon target selection. Conversely, if the U1-U5 motor neurons extend their axons sequentially, then both models remain possible.

To determine the time of U1-U5 axon outgrowth, we used MultiColorFlpOut (MCFO) (Nern et al., 2015) which produces randomized multi-color labeling of neurons within the expression domain of any Gal4 line. We restricted labeling to the NB7-1 lineage using a new split-gal4 killer zipper line (NB7-1-Gal4^KZ^). This new line is based on our published NB7-1-Gal4 line (*ac-VP16 gsb-DBD*) (Kohwi and Doe, 2013) but also includes an R25A05-KillerZipper construct to block expression in NB6-1, which was commonly observed in the previously described NB7-1 split Gal4 line (Kohwi and Doe, 2013; see methods for quantification). This new NB7-1-Gal4^KZ^ line was used for all MCFO or Hb misexpression experiments. Within the NB7-1 lineage, early-born neurons are located close to the midline and later-born neurons are located more laterally (Pearson and Doe, 2003) (Supplemental Fig. 1A). As expected, MCFO labeling of the entire wild-type NB7-1 lineage shows neurons spread from medial to lateral within the CNS, with ipsilateral motor projections and contralateral dendrite projections (Fig. 2A); we call these dendrites because they have a large number of post-synaptic densities but no pre-synaptic sites when analyzed by electron microscopy (Supplemental Fig. 2). We analyzed embryos where MCFO differentially labeled early-born and late-born neurons in the NB7-1 lineage at embryonic stages 12-15 (staging according to Hartenstein, 1993). In all cases, the medial early-born neurons invariably extended axons further than the lateral later-born neurons (Fig. 2A-B; n = 10, p<0.0001, two-tailed unpaired t-test). This observation remained consistent at all tested embryonic stages and independent of the position at which the lineage was subdivided along the medial-lateral axis. In all cases, the U neurons showed staggered axon projections that are remain staggered at every stage; they never stall and become synchronized. Furthermore, in every case where MCFO differentially labeled just a pair of neurons, we always found the medial (early-born) neuron had a longer axon projection than the lateral (later-born) neuron (Fig. 2E-F, n=8, p<0.0001, two-tailed paired t-test). We conclude that during wild type embryonic development, the U1-U5 motor neurons project axons sequentially out the nerve root.

**Fig. 2.**
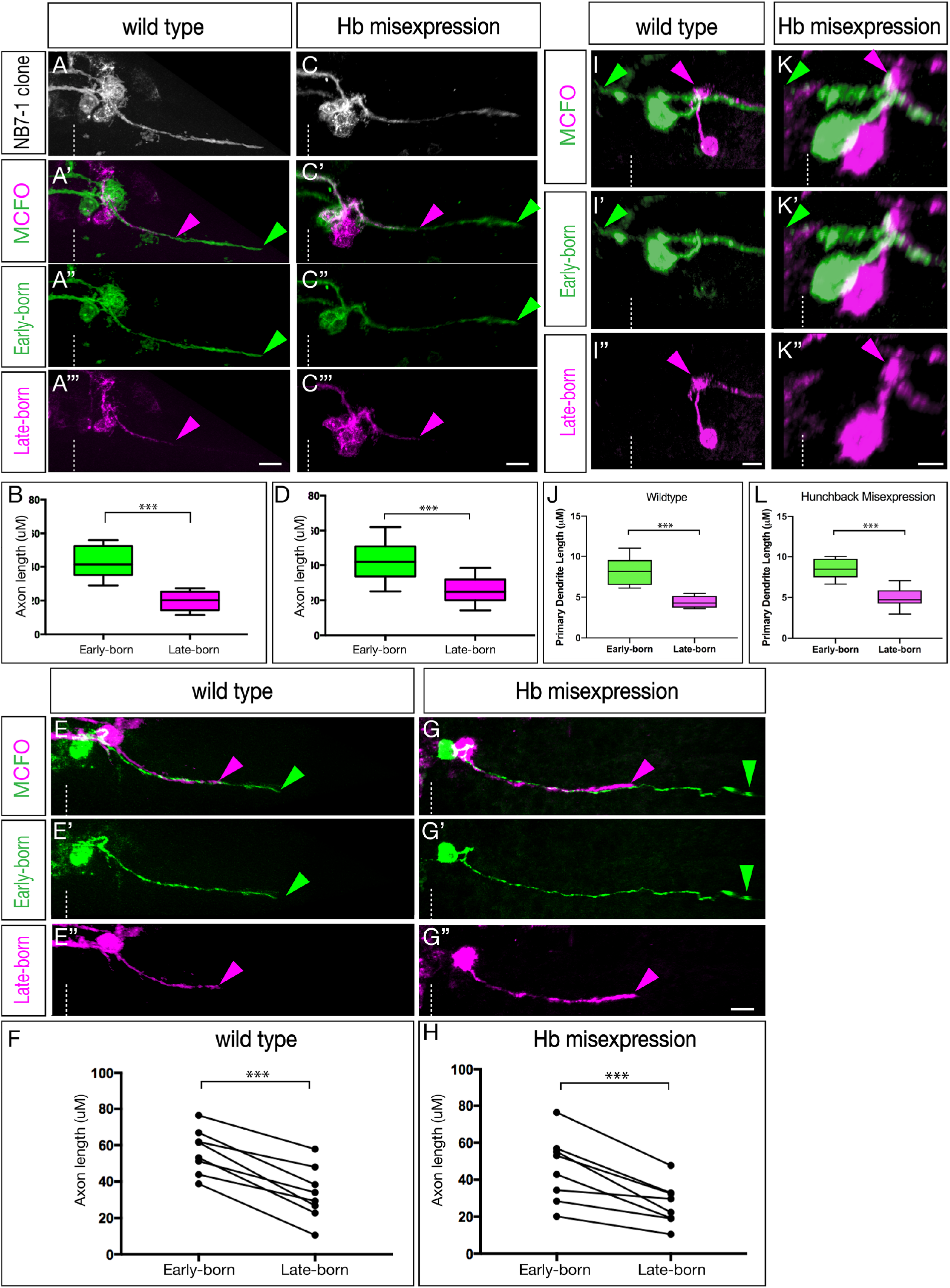
The U1-U5 motor neurons extend axons and dendrites sequentially. (A-H) Axon outgrowth timing in early-and late-born neurons of the NB7-1 lineage in stage 13-15 embryos. (A,B) Wild type multicellular MCFO labeling (*NB7-1-Gal4*^*KZ*^ *UAS-MCFO)*. Analysis was restricted to lineages in which early-born, medially-located, neurons were stochastically labeled in one MCFO color, and the later-born, laterally-located, neurons were stochastically labeled in a different MCFO color. (A) All labeled cells. (A’-A’’’) early-born medial neurons (green) project out of the CNS ahead of later-born lateral neurons (magenta). Arrowheads, most distal axon; dashed line, midline. Scale bar, 5 μM. (B) Quantification of axon length as a representation of timing of axon outgrowth; early-born neurons project further (earlier) than late-born neurons (p < 0.001). (C,D) Hb misexpression (*NB7-1-Gal4*^*KZ*^ *UAS-hb UAS-MCFO*) multicellular MCFO labeling. (C) All labeled cells. (C’-C’’’) early-born medial neurons (green) project out of the CNS ahead of later-born lateral neurons (magenta). Labels and scale bar as in (A). (D) Quantification, as in (B) (p < 0.001). (E,F) Wild type single neuron MCFO labeling (*NB7-1-Gal4*^*KZ*^ *UAS-MCFO)*. (E-E’’) A single early-born medial neuron (green) always projects out of the CNS ahead of a single later-born lateral neuron (magenta). Labels and scale bar as in (A), quantification in (F). (G,H) Hb misexpression (*NB7-1-Gal4*^*KZ*^ *UAS-hb UAS-MCFO*) single neuron MCFO. (G-G’’) A single early-born medial neuron (green) always projects out of the CNS ahead of a single later-born lateral neuron (magenta). Labels and scale bar as in (A), quantification in (H). (I-L) Dendrite outgrowth timing in early-and late-born neurons of the NB7-1 lineage in stage 13-15 embryos. (I,J) Wild type single neuron MCFO labeling (*NB7-1-Gal4*^*KZ*^ *UAS-MCFO)*. A single early-born medial neuron (green) extends a dendrite before a single later-born lateral neuron (magenta). Labels as in (A). Scale bar 5 μM, quantification as in (B) (p < 0.001). (K,L) Hb misexpression (*NB7-1-Gal4*^*KZ*^ *UAS-hb UAS-MCFO*) single neuron MCFO labeling. A single early-born medial neuron (green) extends a dendrite before a single later-born lateral neuron (magenta). Labels as in (I). Scale bar 5 μM, quantification as in (B,D) (p < 0.001).

We next wanted to determine whether misexpression of Hb throughout the NB7-1 lineage, known to produce many ectopic U1 motor neurons (Isshiki et al., 2001; Kohwi and Doe, 2013; Pearson and Doe, 2003), would alter the timing of motor axon outgrowth. We misexpressed Hb in the NB7-1 lineage, and used MCFO to differentially label early-born and late-born neurons. MCFO marking the entire NB7-1 lineage did not change the gross distribution of neurons (Fig. 2C). Importantly, in every case where MCFO differentially labeled early-born and late-born neurons, we found that early-born neurons projected axons out of the CNS before later-born neurons (Fig. 2C-D; n = 10, p<0.0001, two-tailed unpaired t-test). As in the wild-type, in every case where MCFO differentially labeled just a pair of neurons, we always found the more medial (early-born) neuron had a longer axon projection than the lateral (later-born) neuron (Fig. 2G-H, n = 8, p<0.001, two-tailed paired t-test). Moreover, the axon length differential between early-born and late-born neurons was indistinguishable in wild type and Hb misexpression lineages (p<0.001; n= 17, p=0.41, two-tailed unpaired t-test, data not shown).

We next examined the time course of dendrite extension. In wild-type, we observed that earlier-born cells elaborated their dendritic processes before their later-born counterparts (Fig. 2I-J; n = 7, p<0.001, two-tailed unpaired t-test); the same was observed in Hb misexpression animals (Fig. 2K-L; n = 7, p<0.001, two-tailed unpaired t-test). We conclude that sequentially born motor neurons project axons and dendrites sequentially, in both wild type and following Hb misexpression. This raises the question: is intrinsic temporal identity or time of differentiation more important for U1-U5 axon or dendrite target selection?

### Late-born neurons with early intrinsic temporal identity have contralateral dendrite projections

To determine if neuronal morphology was correlated with intrinsic temporal identity or time of differentiation, we first needed to define the morphology of the U1-U5 motor neurons. Previous work has mapped generic U neuron axonal projections (Landgraf et al., 1997), but did not identify muscle targets for specific U1-U5 motor neurons. To precisely define U1-U5 motor neuron identity, we used the serial section transmission electron microscopy (EM) (Ohyama et al., 2015) to reconstruct U1-U5 morphology (Fig. 3A-E; Supplemental Fig. 2). We detected two striking differences in morphology between early-born U1-U2 neurons and later-born U3-U5 neurons: U1-U2 have bipolar projections, whereas U3-U5 have monopolar projections; and U1-U2 have contralateral dendrites, whereas U3-U5 have ipsilateral dendrites (Fig. 3A-E).

**Fig. 3.**
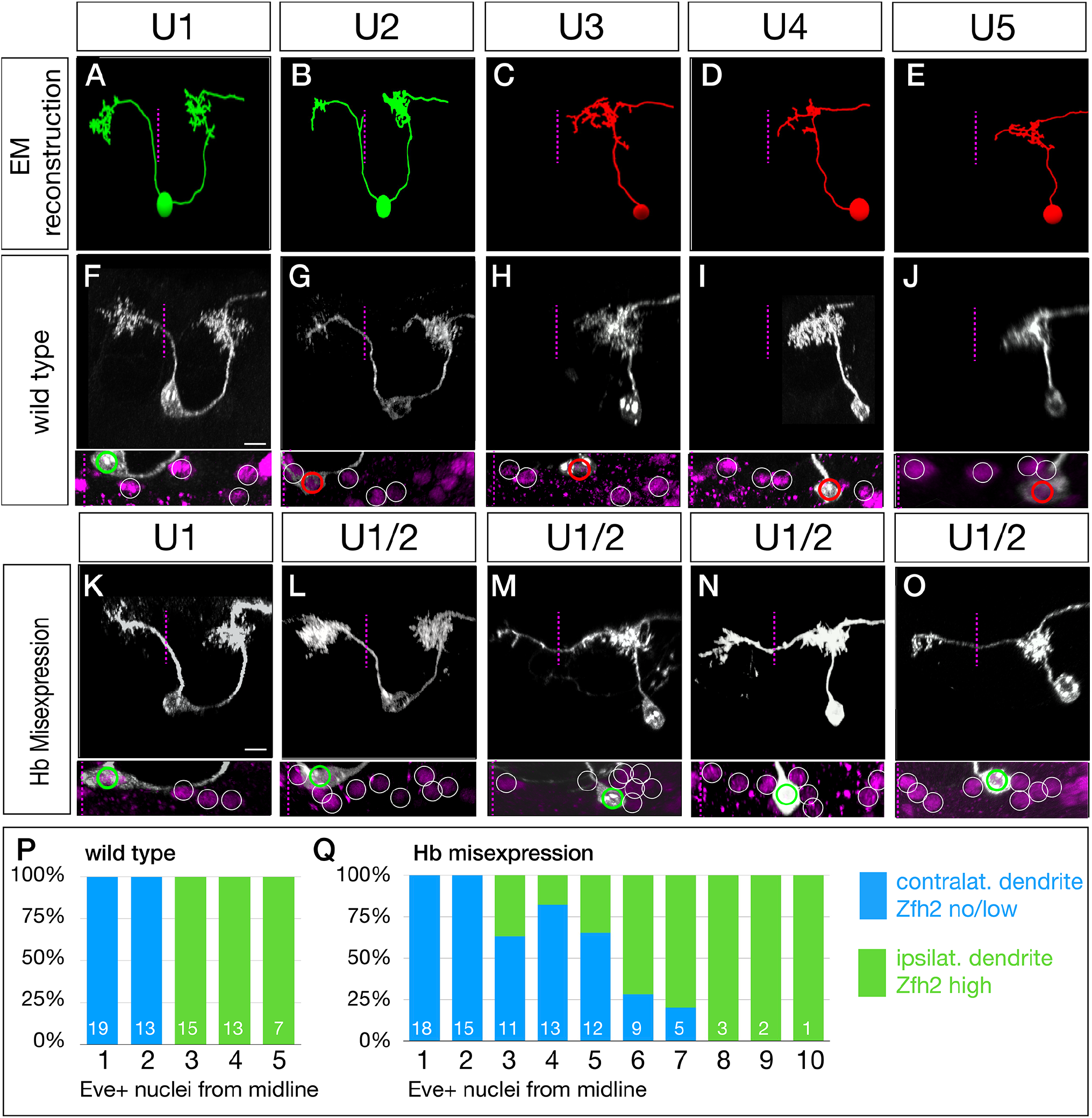
Late-born neurons with early intrinsic temporal identity have “early” dendrite morphology. (A-E) U1-U5 neuronal morphology determined by EM reconstruction in the first instar larval CNS. Early-born U1-U2 neurons (green) have a bipolar morphology with a contralateral dendrite arbor (left of dashed midline), whereas later-born U3-U5 neurons (red) have a monopolar morphology and ipsilateral dendritic arbors. Neuronal birth-order is determined by mediolateral position (U1 most medial/earliest, U5 most lateral/latest). (F-J) Wild type U1-U5 single neuronal morphology by MCFO in L1 larvae (*NB7-1-Gal4*^*KZ*^ *UAS-MCFO*). Neurons are shown from left-right based on birth-order, determined by their position within the five Eve+ neurons (bottom inset). Scale bar, 5 μM. (K-O) Hb misexpression U1-U5 single neuronal morphology by MCFO in L1 larvae (*NB7-1-Gal4*^*KZ*^ *UAS-MCFO*). Neurons are shown from left-right based on birth-order, determined by their position within the Eve+ neurons (bottom inset). The later-born neurons (“ectopic U1”) have acquired a contralateral dendrite, more consistent with their early intrinsic temporal identity than their late time of differentiation. Scale bar, 5 μM. (P,Q) Quantification. In wild type, early-born U1-U2 neurons have low/no nuclear Zfh2, a marker for their early intrinsic temporal identity, and contralateral dendrites; later-born neurons have high Zfh2 and no contralateral projection. In Hb misexpression embryos, all neurons with low/no Zfh2 have a contralateral dendrite, even when they have a late-born time of differentiation (>3 Eve+ nuclei from the midline). The number of neurons scored is shown within each bar.

To determine if the U1-U5 morphology seen in the larval EM reconstruction are reproducible and present in late embryos, we generated MCFO labeling of single U1-U5 motor neurons in late embryos (Fig. 3F-J). Previous work showed U1-U5 stain for the Even-skipped (Eve) transcription factor, and are arranged from medial (U1) to lateral (U5) (Isshiki et al., 2001; Kohwi and Doe, 2013; Pearson and Doe, 2003), which we confirm here (Fig. 3F-J, bottom panels, and quantified in Fig. 3P). We observed that the embryonic U1-U5 motor neurons had a morphology closely matching the larval U1-U5 motor neurons in the EM reconstruction (compare Fig. 3A-E with Fig. 3F-J). We conclude that the early-born U1-U2 neurons and the late-born U3-U5 neurons have distinctive, stereotyped neuronal morphologies.

In wild type, intrinsic temporal identity and time of axon outgrowth are tightly linked; neurons with early intrinsic temporal identity extend axons first, neurons with late intrinsic temporal identity extend axons later. We sought to break this correlation by misexpressing Hb in the NB7-1 lineage so that both early-differentiating and late-differentiating neurons have an early U1 intrinsic temporal identity (Isshiki et al., 2001; Kohwi and Doe, 2013; Pearson and Doe, 2003). To perform this experiment, we needed to monitor neuronal birth-order (a proxy for time of differentiation), intrinsic temporal identity, and neuronal morphology. Birth-order was determined by the neuron position in the medio-lateral series of Eve+ nuclei (medial = early-differentiating; lateral = late-differentiating); intrinsic temporal identity was determined by molecular markers (U1 is Eve+ Zfh2-whereas neurons with later temporal identities are Eve+ Zfh2+; and neuronal morphology was determined by MCFO (Fig. 3K-O). As expected, misexpression of Hb had no effect on the morphology of the endogenous Hb+ U1 or U2 neurons (Fig. 3K,L; U1 n=13). In contrast, all late-differentiating neurons with an ectopic U1 intrinsic temporal identity (Eve+ Zfh2-) had a morphology similar the endogenous U1 neurons: both producing a dorsal, contralateral dendritic arbor (Fig. 3M-O, arrowheads). The penetrance of the transformation declined in neurons with progressively later birthdates (quantified in Fig. 3Q). The failure to project a contralateral dendrite was perfectly correlated with the failure to repress Zfh2 (Fig. 3Q; Supplemental Figure 3), leading us to conclude that these Zfh2+ late-born neurons are simply not being transformed to a U1 identity, and thus fail to project contralaterally. Interestingly, even the transformed ectopic U1 neurons (Eve+ Zfh2-) had their contralateral process emerging from a dorsal location, rather than from the cell body as observed for endogenous U1 neurons, indicating that the morphological transformation was not complete (Fig. 3M-O, Supplemental Movies A-B). Nevertheless, the ectopic U1 neurons are more similar to the endogenous early-born U1-U2 neurons than to the later-born U3-U5 neurons. We conclude that neuronal morphology is more tightly linked to intrinsic temporal identity than to neuronal birth-order.

### Late-born neurons with early intrinsic temporal identity target their dendrites to the U1 dendritic domain

The experiments described above show that late-differentiating neurons with early intrinsic temporal identity have gross morphological features matching their intrinsic temporal identity, rather than their time of differentiation. In this section and the next, we investigate whether these ectopic U1 neurons target their axons to the normal U1 muscle target (the dorsal DO1 and DO2 muscles) and target their dendrites to the normal U1 neuropil target (a contralateral, dorsal volume of neuropil). In this section we assay dendritic projections; in the following section we assay axonal projections.

In wild-type, the endogenous U1 neurons have ipsilateral and contralateral dendrites that are co-localized in the same region of dorsal neuropil, as seen by EM reconstruction (Fig. 4A) or dual color MCFO labeling (Fig. 4B). To map dendrite targeting of “heterochronic” ectopic U1 motor neurons, we misexpressed Hb in the NB7-1 lineage and then screened for MCFO labeling in which one hemisegment had the endogenous U1 motor neuron labeled (identified by its medial position and “U” shaped neuronal morphology), and the opposite hemisegment had an ectopic U1 neuron labeled (identified by its lateral cell body position and dorsal contralateral dendrite process). In every case, we found the “heterochronic” ectopic U1 neuron dendrite precisely targeted to the normal dorsal neuropil target of the U1 neuron, with both ectopic and endogenous U1 dendritic arbors tightly intermingled (Fig. 4B,B’’’; n=10). For each dendrite assessed, correct neuropil localization was confirmed through quantification of the distance of the dendrite from the midline, the anteroposterior distance of the dendrite from the directly anterior hemisegment in relation to the labelled cell, and the position of the dendrite in the dorsoventral axis (Fig. 4C). We observed no significant differences between the dendritic localization of the wild-type U1 neurons and our ectopic early-born cells across any of our positional metrics (Fig.4C, two-way ANOVA, p=0.71). We conclude that intrinsic temporal identity, not time of dendrite outgrowth, generates precise dendrite targeting to the appropriate region of the neuropil.

**Fig. 4.**
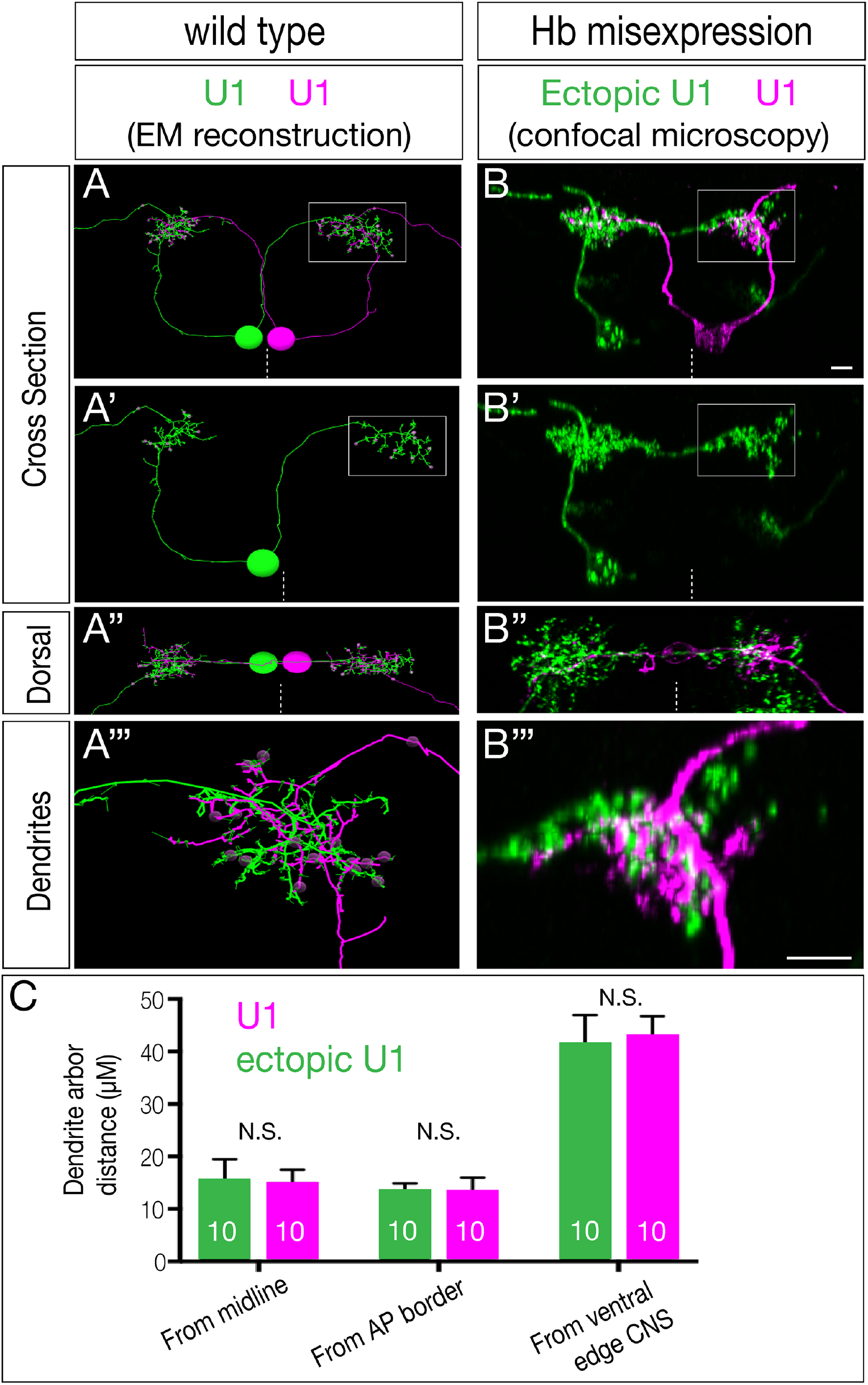
Ectopic U1 dendrites target the normal U1 neuropil domain. (A-A’’’) Wild type bilateral U1 neurons (green, magenta) assayed in the EM reconstruction of the L1 larval CNS. A U1 neuron (magenta) targets its contralateral dendrite to the same neuropil volume as the ipsilateral dendrite of the contralateral U1 neuron (green; boxed region in A,A’ shown enlarged in A’’’). (B-B’’’) Hb misexpression (*NB7-1-Gal4*^*KZ*^ *UAS-hb UAS-MCFO*) assayed by MCFO labeling in L1 larvae, showing an endogenous U1 (magenta; defined by its medial cell body position, bipolar morphology, and contralateral projection), and an ectopic U1 neuron (green; defined by its lateral cell body position, monopolar morphology, and contralateral projection). Note that the endogenous and ectopic U1 neurons target the same dorsal neuropil domain (boxed in B,B’ shown enlarged in B’’’). Midline, dashed line; all views cross section, dorsal up except A’’ and B’’ which are dorsal views, anterior up. Scale bar, 5 μM. (C) Quantification. Endogenous and ectopic U1 dendrites are the same distance from the midline, anterior-posterior (AP) border, and ventral edge of the CNS. n = 10 for U1; n = 10 for ectopic U1.

### Ectopic U1 axons project to dorsal body wall muscles, and lack ventral muscle targets

Here we determine if late-born neurons with early intrinsic temporal identity project their axons to dorsal muscles normally targeted by neurons with early temporal identify, or more ventral muscles normally targeted by late-born neurons. We focus our analysis on L1 larvae, where neuromuscular junctions have formed and are functional for locomotion. In wild-type, we find that the U1-U2 motor neurons innervate the dorsal-most oblique muscles DO1and DO2, and the U3-U5 motor neurons innervate more ventral muscles in the area of DA3 and D04 (Fig. 5A-A’’’; quantified in Fig. 5A’’’’). All motor neurons make varicosities indicating presynaptic differentiation at their muscle targets, but here we do not assay functional synaptic connectivity, simply axon targeting. In contrast, Hb misexpression results in a complete loss of the more ventral axon varicosities, while still exhibiting varicosities at the site of the DO1 and DO2 dorsal muscles (Fig. 5B-B’’’; quantified in Fig. 5A’’’’). These results suggest that later-born motor neurons in the lineage have been transformed into an early intrinsic temporal identity and thereby target the normal early U1-U2 muscle targets.

**Fig. 5.**
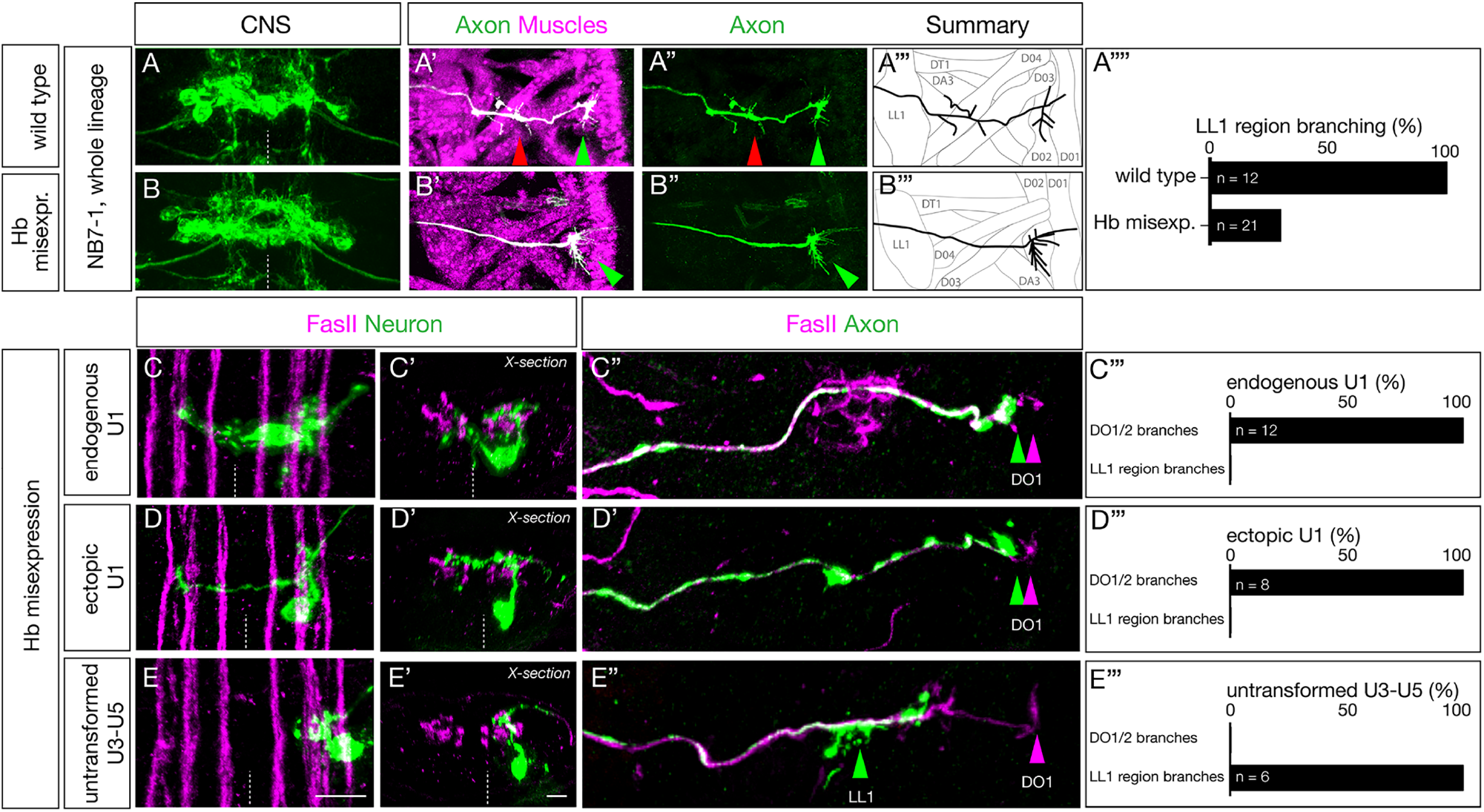
Ectopic U1 axons project to dorsal muscles, and lack ventral muscle targets. (A-B) Wild type (*NB7-1-Gal4*^*KZ*^ *UAS-GFP*) and Hb misexpression (*NB7-1-Gal4*^*KZ*^ *UAS-hb UAS-GFP*) L1 larvae stained for U motor neurons (green) and muscles (magenta). (A-A’’’) Wild type: U motor neurons project axons to dorsal muscles (DO1/DO3) and more ventral muscles in the LL1/DA3 region. (B-B’’’) Hb misexpression: U motor neurons project only to dorsal muscle targets (magenta, diagrammed in B’’’) consistent with ectopic U1-U2 identity at the expense of U3-U5 neuronal identity. (C-E) Hb misexpression L1 larvae (*NB7-1-Gal4*^*KZ*^ *UAS-hb UAS-MCFO*) showing MCFO labeled single neurons. (C-C’’’) The endogenous U1 motor neuron (green; closest to midline). (C) Dorsal view showing medial cell body position, contralateral dendrites, and ipsilateral axon (Zfh2-negative; not shown). (C’) Cross sectional view of same U1 neuron. (C’’) dorsal view of the body wall showing the U1 axon (green arrowhead) projecting to the most dorsal extent of the FasII+ motor neurons (magenta arrowhead). (C’’’) Quantification. (D-D’’’) An ectopic U1 motor neuron (green). (D) Dorsal view showing lateral cell body position, contralateral dendrites, and ipsilateral axon (Zfh2-negative; not shown). (D’) Cross sectional view of the same neuron; note the dorsal origin of the contralateral dendrite and lack of bipolar morphology. (D’’) dorsal view of body wall showing the ectopic U1 axon (green arrowhead) projecting to the most dorsal extent of the FasII+ motor neurons (magenta arrowhead). (D’’’) Quantification. (E-E’’’) A late-born laterally-positioned U3, U4, or U5 motor neuron that was not transformed (based on being Zfh2+; not shown). (E) Dorsal view showing far lateral cell body position, ipsilateral dendrites, and ipsilateral axon. (E’) Cross sectional view of the same neuron. (E’’) dorsal view of the body wall showing the U3-U5 axon (green arrowhead) projecting to a more ventral region along the FasII+ motor neurons (magenta arrowhead). (E’’’) Quantification.

To examine individual motor neurons, we used MCFO following Hb misexpression throughout the NB7-1 lineage. We can observe endogenous early-born U1 motor neurons, identified by their medial position and bipolar morphology (Fig. 5C-C’), that project to DO1/DO2 dorsal muscles (Fig. 5C’’; quantified in Fig. 5C’’’). We also observe “ectopic U1” motor neurons identified by their displacement from the midline, their monopolar morphology, and their dorsal originating contralateral dendrite projection (Fig. 5D-D’); these neurons project to the same DO1/DO2 dorsal muscles as the endogenous U1 motor neuron (Fig. 5D’’; quantified in Fig. 5D’’’). These heterochronic “ectopic U1” motor neurons are clearly different than the normal late-born U3-U5 motor neurons, identified by their lateral position and lack of contralateral dendrites (Fig. 5E-E’), that project to the region of the more ventral muscle LL1 (Fig. 5E’’; quantified in Fig. E’’’). We conclude that intrinsic temporal identity, not time of axon outgrowth, generates precise axon targeting to the appropriate body wall muscles.

To examine the ability of these “ectopic U1” motor neurons to create functional synaptic inputs onto the DO1/DO2 dorsal muscles, we quantified the numbers of pre-synaptic Bruchpilot (Brp) puncta formed by U neurons on their dorsal longitudinal muscle targets. In wild type, U1-U2 neurons form Brp+ synapses on the most dorsal longitudinal muscles DO1/DO2, and the later-born U3-U5 neurons form synapses with the slightly more ventral muscles in the LL1 region (Fig. 6A-D; quantified in Fig. 6I-K). In contrast, the “ectopic U1” motor neurons have a significant shift towards more dorsal muscle targets (Fig. 6E-H). There is a significant loss of pre-synaptic Brp+ puncta in the LL1 region (Fig. 6F, quantified in Fig. 6I), while simultaneously increasing their amount of synaptic input onto the dorsal muscle DO2 (Fig. 6G,H, quantified in Fig. 6J). We saw an insignificant difference in synaptic input onto DO1 between wild type and Hb misexpression (Fig. 6H quantified in Fig. 6K). We conclude that intrinsic temporal identity, not time of axon outgrowth, determines the position of Brp+ presynaptic puncta on the dorsal longitudinal muscle targets.

**Fig. 6.**
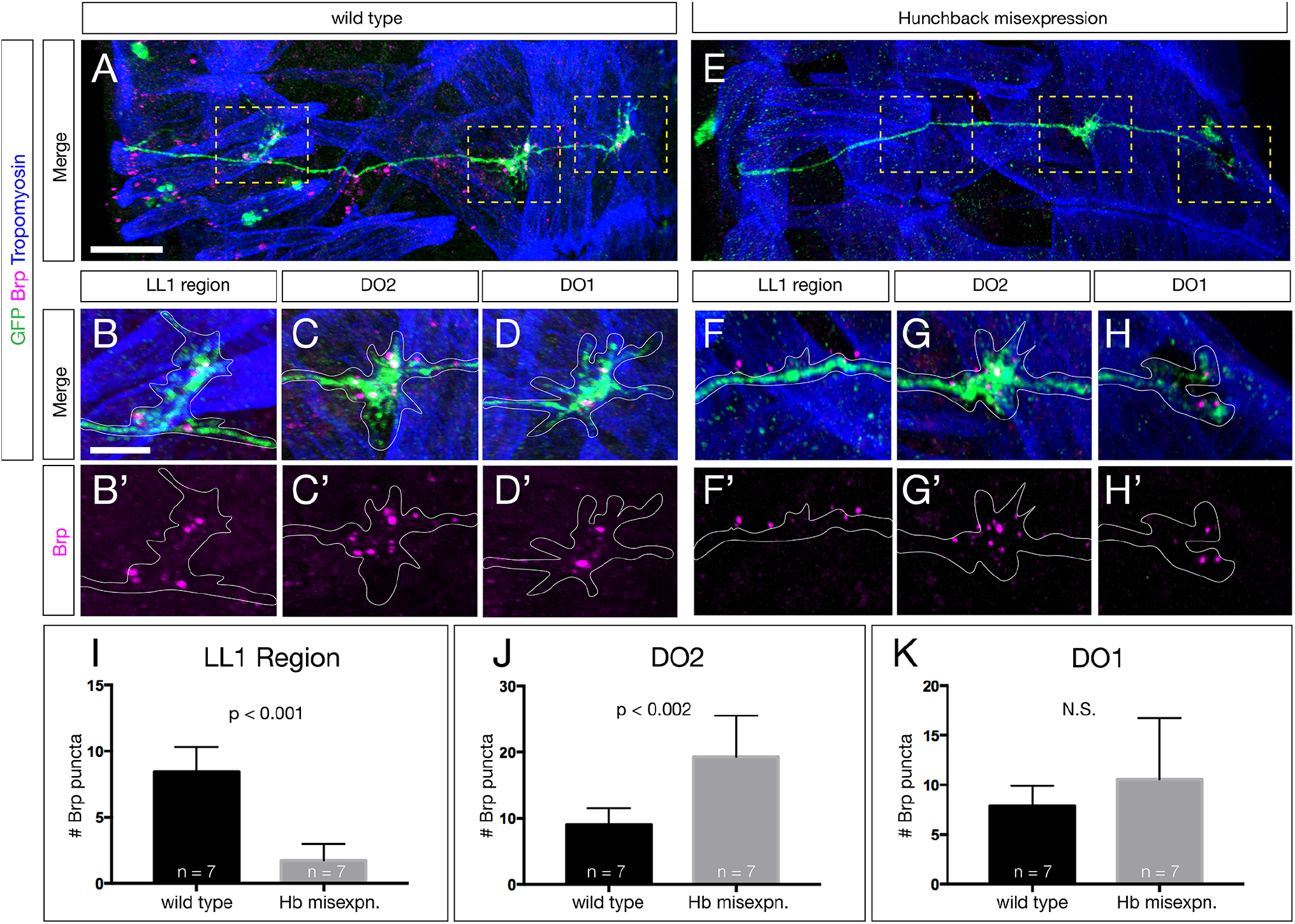
Ectopic U1 axons shift synaptic input from ventral to dorsal muscle targets. (A-D) Wild type L1 larva stained for all U motor neurons in the NB7-1 lineage (GFP, green), Brp+ puncta (magenta) and body wall muscles (Tropomyosin, blue). The U motor neurons have Brp+ puncta contacting muscles around LL1 (B,B’), the DO2 muscle (C,C’) and the DO1 muscle (D,D’). (E-H) Hb misexpression L1 larva (*NB7-1-Gal4*^*KZ*^ *UAS-hb*) stained for all U motor neurons in the NB7-1 lineage (GFP, green), Brp+ puncta (magenta) and body wall muscles (Tropomyosin, blue). There are reduced Brp+ puncta around LL1 (F,F’), increased Brp+ puncta on the DO2 muscle (G,G’) and similar number of Brp+ puncta on the DO1 muscle (H,H’). (I-K) Quantification. Scale bars 15 μm (A,E) and 5 μm (remaining panels).

## Discussion

During neurogenesis, intrinsic temporal identity and time of differentiation are typically tightly correlated. For example, the *Drosophila* NB7-1 sequentially generates the U1-U5 motor neurons which have distinct intrinsic temporal identities and distinct times of differentiation. Our work shows that intrinsic temporal identity is more important than the time of neuronal differentiation for establishing proper axon and dendrite targeting. We generated ectopic motor neurons with an early-born U1 intrinsic temporal identity in a later extracellular environment, breaking the correlation between intrinsic temporal identity and time of differentiation. These late-born ectopic U1 neurons sent their axons to the DO1/2 muscles (together with endogenous U1 neurons), and their dendrites to a dorsal, contralateral neuropil domain (together with endogenous U1-U2 neurons). Furthermore, ectopic U1 neurons are also born in a much more lateral location in the CNS, and yet are able to find their correct axon and dendrite targets. Overexpression of Hunchback in other neuroblast lineages generates early-born neuronal identity based on molecular marker expression (Isshiki et al., 2001; Moris-Sanz et al., 2015; Novotny et al., 2002; Tran and Doe, 2008), but here we characterize the pre-and post-synaptic targeting of these “heterochronic” neurons. Our data show that intrinsic temporal identity is an important determinant of neuronal axon and dendrite targeting.

Temporal transcription factors (TTFs) are known to regulate neuronal cell fate in multiple neuroblast lineages in *Drosophila* (Doe, 2017). In mushroom body neuroblasts, TTFs are known to specify the molecular and morphological features of the Kenyon cells (Kao and Lee, 2010; Liu et al., 2015; Zhu et al., 2006). Similarly, in type II neuroblast intermediate neural progenitor (INP) lineages, recent work has shown that the late TTF Eyeless specifies the molecular identity and axon/dendrite targeting of several classes of central complex neurons (Sullivan et al., 2019). In contrast, the INP parental type II neuroblasts express a different set of TTFs, but nothing is yet known about their role in axon/dendrite targeting (Ren et al., 2017; Syed et al., 2017). Recent work has shown that optic lobe neuroblasts express TTFs that specify the molecular identity and axon targeting of visual system neuronal subtypes, but it is unknown if sequentially born neurons project axons sequentially or synchronously (Bertet et al., 2014; Erclik et al., 2017; Li et al., 2013b). The antero-dorsal larval brain neuroblast expresses TTFs that govern the identity of olfactory projection neurons, as well as regulating the dendritic targeting to specific antennal lobe glomeruli, but it is unknown if the projection neuron dendrites project sequentially or synchronously (Jefferis et al., 2001; Liu et al., 2015). Another relevant system for studying temporal identity and axon targeting is the sequential production of R8 photoreceptor neurons, which express a graded level of the transcription factor Sequoia based on their birth-order; abolishing the Sequoia gradient prevents the smooth distribution of R8 axon terminals in the medulla neuropil (Kulkarni et al., 2016; Petrovic and Hummel, 2008). Taken together, abundant data suggest that TTFs control neuronal molecular identity, with a growing number of studies showing that TTFs also regulate axon/dendrite targeting to specific neuropil domains or muscles.

Previous work has demonstrated that motor neurons in different lineages project axons at different times, e.g. aCC prior to RP2 (Sanchez-Soriano and Prokop, 2005). Similarly, we found that the U1-U5 motor neurons extend axons sequentially, and independently of their intrinsic temporal identity. This suggests that the initial timing of axon extension is regulated by an internal clock mechanism in each cell, likely beginning upon its terminal cell division. In *C. elegans*, the HSN motor neurons require expression of *lin-18* mRNA to initiate axon extension (Olsson-Carter and Slack, 2010); whether a similar mechanism is used by U1-U5 motor neurons is unknown. We also show that dendrite elaboration occurs much later than axon extension in the U motor neurons. The observation that axon outgrowth precedes dendrite outgrowth has been widely reported (Gerhard et al., 2017; Mason, 1983; Mumm et al., 2006; Ramon y Cajal, 1909), although the mechanism setting the time of axon or dendrite outgrowth is poorly understood.

Hb misexpression robustly transformed later-born U motor neurons into ectopic U1 neurons, yet there were two limitations. First, ectopic U1 neurons do not have a bipolar cell body; they branched off dendrites from the dorsal axon, rather than from the cell body. Nevertheless, despite their novel dorsal outgrowth, ectopic U1 dendrites targeted a contralateral neuropil volume indistinguishable from the endogenous U1 neurons. The failure of the ectopic U1 neurons to generate a bipolar somata may be due to (i) incomplete transformation of neuronal identity, (ii) an abnormal lateral cell body position, (iii) changing extrinsic cues, or (iv) intrinsic changes in the neuronal cytoskeletal that are not under Hb regulation. A second limitation is the decline in Hb potency as the NB7-1 lineage progresses. We find that following Hb misexpression there are always some laterally-positioned Eve+ motor neurons that fail to repress Zfh2 and fail to extend contralateral dendrites (Supplemental Figure 3); we conclude these neurons are simply untransformed. The inability of Hb to fully transform late-born neurons has been well documented (Kohwi et al., 2013; Pearson and Doe, 2003). The striking correlation between Zfh2 expression and ipsilateral dendrite projection raises the possibility that Zhf2 levels regulate dendrite midline crossing. Testing this hypothesis would require generating *UAS-zfh2* transgenics for NB7-1-specific overexpression, and an FRT *zfh2* mutant fourth chromosome to make *zfh2* mutant clones in NB7-1.

The NB7-1 cell cycle is ∼50 min (Hartenstein et al., 1987) which means that ectopic U1 motor neurons can be born up to 6 divisions or 300 min later than normal and yet still find their normal axon and dendrite targets. This suggests that the guidance cues used for endogenous U1 pathfinding are still present many hours later. Consistent with this, the major pathfinding ligands regulating sensory axon targeting and motor dendrite targeting in the CNS – NetrinA/B, Slit, Semaphorin1/2, and Wnt5 (Mauss et al., 2009; Wu et al., 2011; Yoshikawa et al., 2016; Zlatic et al., 2003; Zlatic et al., 2009) – all maintain their graded expression patterns during this window of neurogenesis (Fradkin et al., 2004; Harris et al., 1996; Mitchell et al., 1996; Rothberg et al., 1988; Yoshikawa et al., 2003; Zlatic et al., 2009). Although we can’t exclude the possibility of the endogenous and ectopic U1 neurons using different cues to find their proper targets, e.g. later-born neurons may project along “pioneer neuron” processes formed earlier in neurogenesis, it is more likely that both early-born endogenous U1 neurons and later-born ectopic U1 neurons use the same guidance cues for axon and dendrite targeting.

In the future, it will be important to understand the mechanism by which the Hb transcription factor confers U1 neuron axon and dendrite targeting. As mentioned above, it is likely that the endogenous and ectopic U1 motor dendrites target the proper neuropil domain by responding to the known Netrin, Slit, Semaphorin and Wnt5 ligand gradients (Mauss et al., 2009; Wu et al., 2011; Yoshikawa et al., 2016; Zlatic et al., 2003; Zlatic et al., 2009). Thus, we hypothesize that Hb induces expression of distinct receptor combinations that allow the endogenous and ectopic U1 axon and dendrite to respond to these persistent pathfinding ligand gradients. Hb may directly regulate receptor gene expression, or it may act via an intermediate tier of transcription factors, similar to the “morphology transcription factors” that act downstream of temporal transcription factors in establishing adult leg motor neuron axon and dendrite targeting (Enriquez et al., 2015). Understanding how Hb directs axon and dendrite targeting will require characterization of receptor expression in endogenous and ectopic U1 neurons, and/or single cell RNA-seq to characterize the endogenous and ectopic U1 neuron transcriptomes.

## MATERIALS AND METHODS

### Fly Stocks

Male and female *Drosophila melanogaster* were used. The chromosomes and insertion sites of transgenes (if known) are shown next to genotypes. Previously published Gal4 lines, mutants and reporters used were: *hs-FLPG5;;MCFO* (I and III; FBst0064086), *UAS-hunchback*/*CyO* (II) (Isshiki et al., 2001), *10XUAS-IVS-mCD8::GFP* (III, FBst0032185).

### New NB7-1-Gal4^KZ^ line

We generated a new NB7-1-Gal4^KZ^ line that uses an enhancer killer zipper construct to eliminate the NB6-1 off-target expression seen in the published NB7-1 split-Gal4 line (Kohwi et al., 2013). The previous NB7-1-Gal4 line showed NB6-1 expression in 65% of hemisegments (n=20); the NB7-1-Gal4^KZ^ line shows NB6-1 expression in just 25% of hemisegments (n=20). The new split gal4 genotype is *ac-VP16 gsb-DBD, R25A05-KillerZipper/CyO* (II; attP40), where R25A05 is an enhancer expressed in NB6-1. The full stock was created as follows. Syn21-KZip(+)-P10 fragment was PCR amplified from CCAP-IVS-Syn21-KZip(+)-P10 (Dolan et al., 2017) a gift from Benjamin White (NIH/NIMH) and fused via Gibson assembly with NheI/HindIII digested pBPGal80Uw-6 (Pfeiffer et al., 2008) to create pBP-Syn21-KZip. The Janelia Research Campus enhancer R25A05 (FBst0000162964) was introduced into pBP-Syn21-KZip by gateway cloning (Pfeiffer et al., 2008) to generate *R25A05-KZip*, which was then integrated into attP40 site by standard injection (Bestgene Inc. Chino Hills, CA).

### Immunofluorescence staining

Primary antibodies were: rabbit anti-Hunchback #5-25 (1:200) (Tran and Doe, 2008), rabbit anti-Eve #2472 (1:100, Doe Lab), chicken anti-GFP (1:1000, Abcam, 13970), rat anti-Zfh2 (1:250) (Tran et al., 2010), mouse anti-HA-Alexa Fluor 488 conjugate (1:200, Cell Signaling, 2350S), rat anti-HA (1:100, Sigma #11867423001), chicken anti-V5 (1:1000, Bethyl, A190-218A), rat anti-FLAG (1:400, Novus, NBP1-06712), and mouse anti-FasII (1:60, DSHB, 1D4). Secondary antibodies were from Molecular Probes or Jackson ImmunoResearch and were used at 1:350.

Embryos were blocked overnight in 0.3% PBST (1X PBS with 0.3% Triton X-100) with 5% normal goat serum and 5% donkey serum (PDGS), followed by incubation in primary antibody overnight at 4°C. Next, embryos underwent three 30 minute washes in PBST, followed by an overnight secondary antibody incubation at 4°C. Embryos were then dehydrated in a glycerol series (10%, 50%, 90%) for 20 minutes each followed by 90% glycerol with 4% n-propyl Gallate overnight before imaging.

Whole L1 larvae were washed for 2 h in methanol, blocked overnight in 0.3% PBST (1X PBS with 0.3% Triton X-100) with 5% normal goat serum and 5% donkey serum (PDGS), followed by incubation in primary antibody for two nights at 4°C. Next, larvae were washed overnight in PBST, followed by two nights secondary antibody incubation at 4°C. Embryos were dehydrated in a glycerol series (10%, 50%, 90%) for 20 minutes each followed by 90% glycerol with 4% n-propyl Gallate overnight before imaging. Larval brains were dissected in 0.3% PBST, fixed in 4% paraformaldehyde in PBST, rinsed, and blocked in PDGS with 0.3% Triton X-100. Staining was carried out as above for embryos, but after the secondary antibody incubation brains were mounted in Vectashield (Vector Laboratories).

### MCFO labeling

MCFO labeling in wildtype used *ac-gsb-Gal4, R25A05-KillerZipper* (II) *x hsFLPG5;;UAS-MCFO* (I and III) and in Hb misexpression used *ac-gsb-Gal4, R25A05-KillerZipper* (II) *x hsFLPG5;UAS-Hunchback;UAS-MCFO* (I, II and III). Embryos were collected for 2 h at 25°C, aged 4 hours and heat shocked at 37°C (15-20 minutes for dense labeling, 8-10 minutes for sparse labelling), then left to develop until desired stages.

### Imaging

Images were captured with a ZeissLSM 710 or ZeissLSM 800 confocal microscope with a *z*-resolution of 0.35 µm. Images were processed using the open-source software FIJI (https://fiji.sc) and Photoshop (Adobe). Figures were assembled in Illustrator CS5 (Adobe). Three dimensional reconstructions, morphometrics and level adjustments were generated using Imaris (Bitplane). Any level adjustment was applied to the entire image.

### Statistical Analysis

Statistical significance is denoted by asterisks: ****p<0.0001; ***p<0.001; **p<0.01; *p<0.05; n.s., not significant. The following statistical tests were performed: Two-tailed unpaired t-test (Figures 2B,2D,2J,2I,6H,6I,6J); Two-tailed paired t-test (Figs. 2F,2H); and 2-Way ANOVA (Figure 4). All analyses were performed using Prism8 (GraphPad). The results are stated as mean ± s.d., unless otherwise noted.

### Serial section electron microscopy

We accessed a previously published serial section transmission electron microscopic volume of the newly hatched larval CNS using CATMAID software (Ohyama et al., 2015) to describe the U1-U5 motor neurons in the first abdominal segment. U1-U5 motor neurons were identified based on their published unique dendritic morphology (Landgraf et al., 1997).

## Acknowledgements

We thank Luis Sullivan and Judith Eisen for comments on the manuscript; Sen-Lin Lai for making the new NB7-1-Gal4^KZ^ line; Aref Zarin for annotating U3-U5 neurons in the EM reconstruction; Brandon Mark for providing helpful advice through the work, and Cooper Doe for assistance with sample preparation and imaging. Transgenic lines were made by BestGene (Chino Hills, CA). Stocks obtained from the Bloomington *Drosophila* Stock Center (NIH P40OD018537) were used in this study.

## Author contributions

CQD and AQS conceived of the project, AQS performed experiments, and CQD and AQS wrote the paper. Both authors commented and approved of the manuscript.

