## supplemental figures for "The Hunchback temporal transcription factor determines motor neuron axon and dendrite targeting in *Drosophila*"

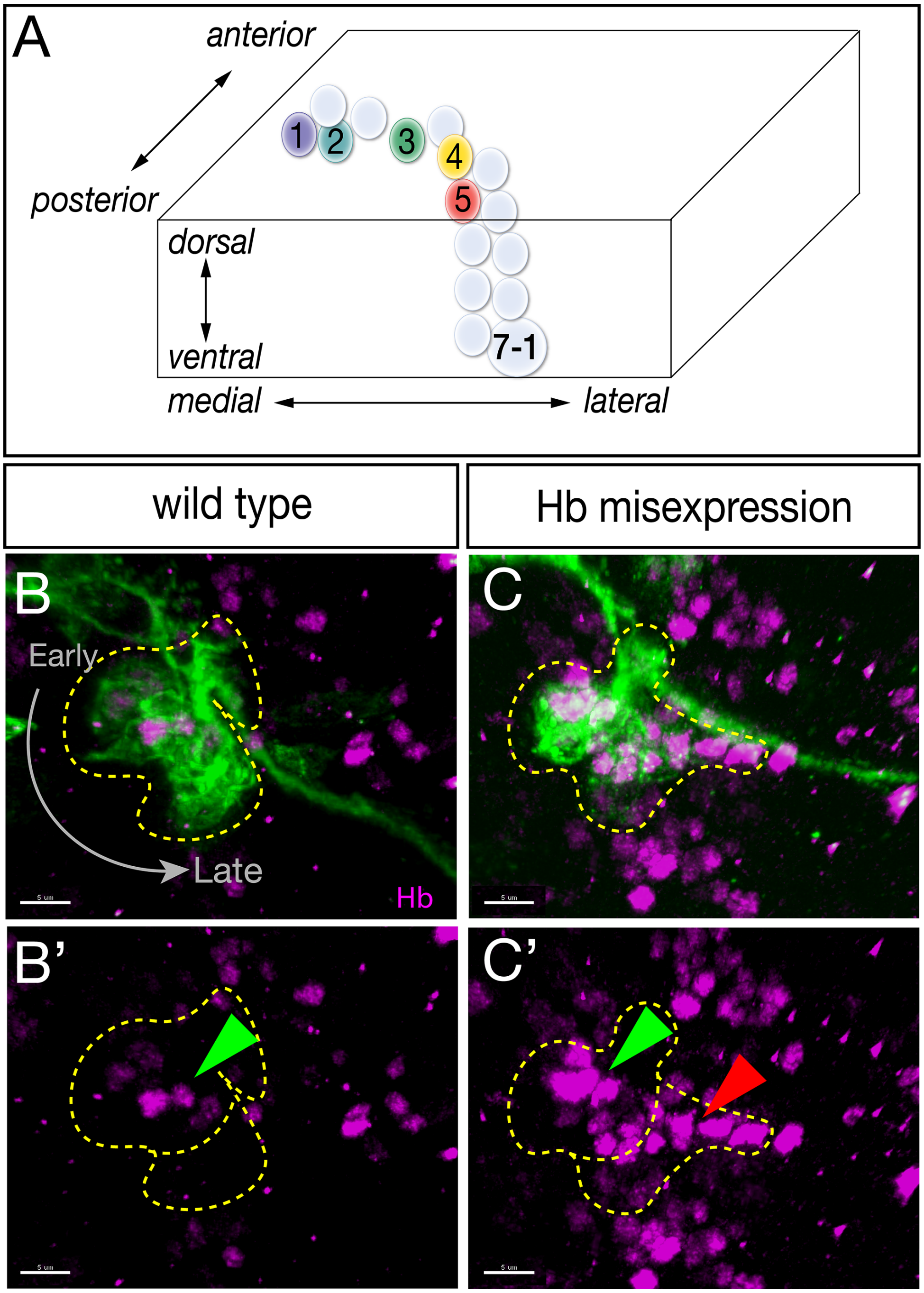


**Supplemental Fig. 1. Introduction to the NB7-1 lineage**

(A) NB7-1 moves laterally as it divides, resulting in early-born neurons (U1-U2) having a medial position and later-born neurons (U3-U5) having a more lateral position (Pearson et al., 2003). (B) In wild type, NB7-1-Gal4 used for dense MCFO shows most or all progeny (green), but only medial cells in the lineage are Hb+ (magenta, green arrowhead); lineage outlined in dashed lines. (C) Hb misexpression (magenta) throughout the NB7-1 lineage results in Hb+ early-born neurons (green arrowhead) and Hb+ late-born neurons (red arrowhead); most of these late-born neurons have an early temporal identity (Kohwi et al., 2013).

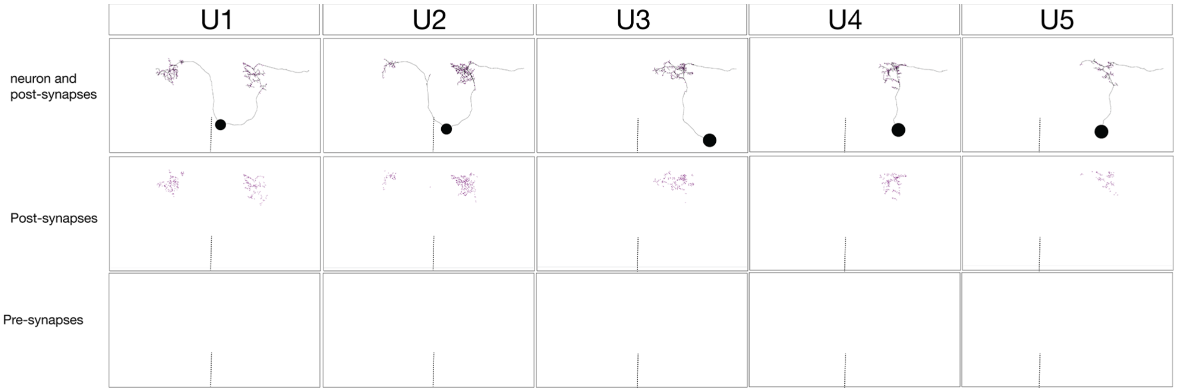


**Supplemental Fig. 2. U1-U5 motor neurons have dendrites within the neuropil that have abundant post-synaptic input but no pre-synaptic output.**
Serial section transmission electron microscopy volume of the entire newly hatched larval CNS contains U1-U5 motor neurons in segment A1 left (black tracing, top row). U1-U5 were identified by their unique neuronal morphology and dendritic arbor location in the neuropil (see Methods). Top row: U1-U5 neurons with post-synapses shown (blue dots). Middle row: the post-synapses are shown (blue dots). Bottom row: the pre-synapses are shown; there are no pre-synapses, and thus we call the CNS arbors “dendritic.” Midline, dashed line.

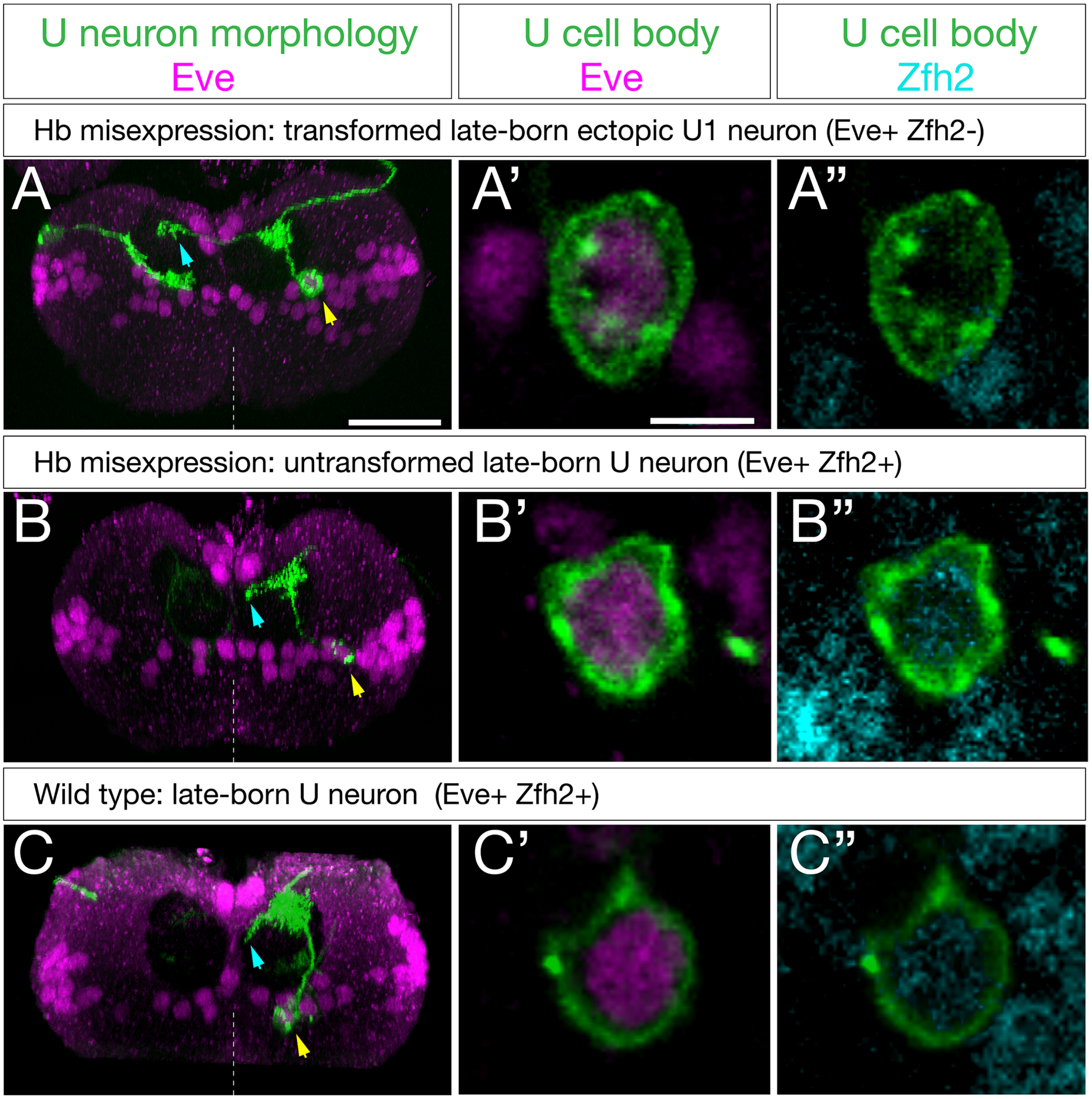


**Supplemental Fig. 3. U neurons that fail to repress Zfh2 fail to project contralaterally.** (A) Representative example of Hb misexpression in the NB7-1 lineage transforming later-born, laterally-positioned U neurons into an ectopic U1 identity based on Zfh2 repression and contralateral dendrite projections. (A', A'') Enlargement of somata shown in (A).
(B) Representative example of Hb misexpression in the NB7-1 lineage failing to transform a later-born, laterally-positioned U neuron based failure to repress Zfh2 and failure to extend a contralateral dendrite projection. (B', B'') Enlargement of somata shown in (B).
(C) Representative example of a wild type later-born, laterally-positioned U3-U5 neuron based Zfh2 expression and failure to extend a contralateral dendrite projection. (C', C'') Enlargement of somata shown in (C).
Posterior view, dorsal up, midline dashed. Scale bars, 15μm (A,B,C), 5 μm (other panels).
